# The Nucleocapsid Protein Of SARS-CoV-2, Combined With ODN-39M, Is A Potential Component For An Intranasal Bivalent Pancorona Vaccine

**DOI:** 10.1101/2022.06.02.494502

**Authors:** Yadira Lobaina, Rong Chen, Edith Suzarte, Panchao Ai, Vivian Huerta, Alexis Musacchio, Ricardo Silva, Changyuan Tan, Alejandro Martin, Laura Lazo, Gerardo Guillén, Ke Yang, Yasser Perera, Lisset Hermida

## Abstract

Despite the rapid development of vaccines and their reported efficacy for controlling the COVID-19 waves, two key challenges remain: the scope of the immunity against upcoming variants and zoonosis events, and the induction of mucosal immunity able to clear the virus in the upper respiratory tract for halting the transmission. The present study is aiming at assessing a potential component for a new generation of vaccines so as to overcome such limitations. The recombinant nucleocapsid (N) protein from SARS-CoV-2 Delta variant was combined with a phosphodiester backbone CpG ODN (ODN-39M), forming high molecular weight aggregates. The evaluation of its immunogenicity in Balb/C mice revealed that only administration by intranasal route induced a systemic cross-reactive Cell-Mediated-Immunity (CMI). In turn, this combination was able to induce anti-N IgA in lungs, which along with the specific IgG in sera and CMI in spleen, resulted cross-reactive against the nucleocapsid protein of SARS-CoV-1. Furthermore, the nasal administration of the N+ODN-39M preparation combined with the RBD Delta protein, as inductor of neutralizing Abs, enhanced the local and systemic immune response against RBD with a modulation toward a Th1 pattern. Taken together, these results make the N+ODN-39M preparation a suitable component for a future intranasal pancorona vaccine against Sarbecoviruses. Particularly, the bivalent vaccine formulation N+ODN-39M+RBD could be used as an effective nasal booster in previously vaccinated population.

## 1. Introduction

The ongoing SARS-CoV-2 pandemic is a public health crisis that has resulted, to date, in the loss of over 6 million lives and caused an unprecedented disruption to humanity. Laboratories across the globe have worked intensively in developing different vaccine against SARS-CoV-2. In general, most of the approved vaccines have been able to induce more than 50% of protection, the minimal limit requested by WHO and have contributed to control the current waves of SARS-CoV-2 variants (Al Kaabi et al., 2021) (Li et al., 2021) (Lopez Bernal et al., 2021) (Mas-Bermejo et al; 2022).

Despite their undoubtedly relevant role they have two common limitations. First, vaccines are based on the spike (S), or RBD (receptor binding domain), proteins from the first circulating clades of SARS-CoV-2 (Walsh et al., 2020); therefore, the immunity induced is mainly directed to the close SARS-CoV-2 variants of the Ancestral one (Andrews et al., 2022). Second, the mucosal immunity is not properly induced; consequently, the viral transmission cannot be halted (Russell et al., 2020).

SARS-CoV-2 infection is the third zoonosis related to human lethal coronaviruses after SARS-CoV in 2002 and MERS-CoV in 2012. Even more alarming, The USAID PREDICT 1 program (2009-2019) identified 113 novel coronaviruses in animals and people in ecological hotspots with intensive spill over interfaces. These facts, along with the appearance of new variants of SARS-CoV-2, has guided to the scientific community to propose a new generation of vaccines with broader scope of protection: the pancorona vaccines (Rubin, 2021) (Morens et al., 2022). So far, two main approaches are being addressed, multiple immunogenic antigens/regions based on S protein (multivalent) and proteins/designs based on conserved regions among coronaviruses. A multivalent approach requires the presence of various immunodominant regions, and the scope is generally limited to the variants of such regions (Cohen et al., 2021; Liang et al., 2021; Wang et al., 2022). Alternatively, the approach based on conserved antigens could reach a broad scope depending on the level of conservation of the antigen and its proper presentation. The nucleocapsid (N) protein is one example of conserved antigen which has been tested in different vaccine platforms and combinations with encouraging results against SARS-CoV-2 (Chiuppesi et al., 2020)(Ahn et al., 2021) (Thura et al., 2021).

On the other hand, the selection of a suitable adjuvant is crucial in the design of new vaccine formulations. CpG ODNs are considered potent enhancers of the immune response since they bind to and activate Toll-like receptor -9 (TLR9) for initiating an important innate immune response. In fact, more than 100 clinical trials using CpG ODNs have been conducted for assessing their use in preventing or treating allergy, infectious diseases and cancer (Scheiermann et al., 2015). In the present work, we combined the recombinant nucleocapsid protein, a conserved antigen from SARS-CoV-2, with the ODN-39M, a synthetic CpG ODN, to favor the formation of aggregate conformations of the N protein based on its natural capacity to bind RNA. This ODN interacts with other viral capsid proteins and exerts adjuvant effect (Gil et al., 2015), (Gil et al., 2016), (Olivera et al., 2020). The resulting preparation was evaluated in mice by different administration routes. Interestingly, the N+ODN-39M preparation, administered by intranasal route, induced the highest anti-N CMI response in spleen and also elicited humoral immunity in both, sera and lungs. Importantly, the overall immunity generated was cross-reactive against N protein from SARS-CoV-2 Delta and Omicron variants, as well as from SARS-CoV-1. Furthermore, a nasal bivalent formulation, based on N+ODN-39M preparation combined with RBD Delta protein, enhanced the local and systemic immune response against RBD with a modulation toward a Th1-like pattern. These findings support the use of the N+ODN-39M preparation as a potential component of a future intranasal pancorona vaccine.

## 2. Materials and Methods

### Recombinant proteins, peptide and ODN-39M

The recombinant antigens were purchased from Sino Biological Inc (China). N proteins from: SARS-CoV-2, Delta variant (40588-V07E29), Omicron variant (40588-V07E34); SARS-CoV-1 (40143-V08B); MERS-CoV (40068-V08B); HCoV-229E (40640-V07E), and RBD proteins from SARS-CoV-2: Delta variant (40592-V08H90), Ancestral variant (40592-VNAH) and ACE-2-His (10108-H08B).

The peptide N_351-365_ from SARS-CoV-2 (ILLNKHIDAYKTFPP) was synthesized with ≥ 97% purity by Zhejiang Peptides Biotech (China).

The ODN-39M, a 39 mer, whole phosphodiester backbone ODN (5’-ATC GAC TCT CGA GCG TTC TCG GGG GAC GAT CGT CGG GGG-3’), was synthesized by Sangon Biotech (China).

### In vitro aggregation procedure of N protein with ODN-39M

The N protein from SARS-CoV-2, Delta variant, was subjected to *in vitro* aggregation, as previously described (Gil et al., 2015), with few modifications. Briefly, in a 100 µL of reaction, 40 µg of N protein were mixed with different quantities of ODN-39M (2 µg, 4 µg, 8 µg, 16 µg, 40 µg, 60 µg and 80 µg) in 10 mM Tris, 6 mM EDTA (pH 6.9) buffer. The mixtures were incubated for 30 min at 30°C in water bath and after, were stored at 4°C for 4 hours. Finally, each preparation was centrifuged at 14 000xg for 10 min. The resulting supernatant was collected and tested for protein concentration. For animal evaluations the mixture of 40 µg of N protein and 60 µg of ODN was selected (mass ratio: 0.66). This preparation was characterized by SDS-PAGE and agarose electrophoresis, in its natural condition and after crosslinking with 0.5 and 1% formaldehyde (FA). After the treatment, FA-treated samples were quenched by adding Tris to a final concentration of 0.1M.

### Immunization experiments

Adult (6 to 8 weeks old) females Balb/c mice (inbred, H-2d) were housed at Beijing Vital River Laboratory Animal Technology Co., Ltd. The standard of laboratory animal room complied with the national standard of the people’s Republic of China GB14925-2010. All the experimental protocols were approved by Institutional Animal Care and Use Committee.

Groups of five to ten animals each were immunized with three doses, administered on days 0, 7 and 21 by intranasal (i.n) or subcutaneous (s.c) routes, according to each experimental design. Formulations for s.c administration were prepared with aluminum hydroxide (alum; Alhydrogel, Invitrogen), as adjuvant, at a final concentration of 1.4 mg/ml. A dose of 10 μg, of each protein per mouse, was evaluated. All the immunogens were dissolved in sterile PBS. For i.n and s.c administrations, the immunogen was administered in a final volume of 50 μL and 100 μL, respectively. In each experiment, placebo immunized groups were included as controls by each route.

In the first mice experiment, six groups of six animals each were employed. Groups 1 and 2 received N protein and N+ODN-39M, respectively, adjuvated with alum by s.c route. Groups 3 and 4 were intranasally immunized with N and N+ODN-39M, respectively. Groups 5 and 6 acted as controls receiving PBS+alum (s.c) and PBS (i.n), respectively. Three animals per group were sacrificed on days 12 and 18 after the last immunization.

For the second mice experiment, three groups of five animals each were used. Groups 1 and 2 were intranasally immunized with N and N+ODN-39M, respectively. Group 3 received PBS (i.n). Animals were sacrificed on day 26 after the last dose.

The third experiment was carried out with six groups of five animals each. All groups were intranasally immunized. Groups 1 and 2 received N and N+ODN-39M, respectively. Groups 4 and 3 received the same formulations than 1 and 2, respectively, but including RBD. Groups 5 and 6 were immunized with RBD and PBS, respectively. Animals were sacrificed 12 days after the last immunization.

At the indicated time points three types of samples were collected: Sera, Bronchoalveolar Fluid (BALF) and spleens.

### Assessment of humoral immune response by ELISA and a surrogate virus neutralization test

The antibody response in sera and BALF was monitored by ELISA. Briefly, anti-IgG, subclasses and, -IgA ELISAs were carried out as previously described (Lobaina et al., 2010). Briefly, 96 well high-binding plates (Costar, USA) were coated with N (3 μg/mL) or RBD (2 μg/mL) proteins and blocked with 2% skim milk solution. Samples were evaluated in duplicates using different dilution starting from 1/50 for sera. BALF were assayed directly, without dilution. Specific horseradish peroxidase conjugates (Sigma, USA) were employed and OPD (Sigma, USA)/hydrogen peroxide substrate solution was used. After 10 min of incubation in the dark, the reaction was stopped using 2 N sulfuric acid and the optical density (O.D) was read at 492nm in a multiplate reader (FilterMax F3, Molecular Devices, USA). For antibody response measured in sera data was represented in the graphics as log10 titers. The arbitrary units of titers were calculated by plotting the O.D values obtained for each sample in a standard curve (a hyper-immune sera of known titer). The positivity cut-off was established as 2 times the average of O.D obtained for a pre-immune sera pool. On the other hand, for antibody response in BALFs, since the positive signals obtained are usually lower, results were represented as O.D at 492 nm.

For detecting functional antibodies generated by RBD formulations, a surrogate virus neutralization test (sVNT) was used that allows to determine the capacity of mice sera to inhibit the interaction of RBD with ACE2 protein. To evaluate the inhibitory activity against the ancestral strain, mice sera were assayed using the SARS-CoV-2 Inhibitor Screening ELISA kit (Sino Biological, KIT001), according to the supplier instructions. In this assay, antibodies present in sera compete with ACE2 for binding to RBD (Ancestral variant) coating the wells of the plate. Bound ACE2 is detected via the polyhistidine tag using an anti-polyhistidine Mab conjugated to HRP. In this experimental set up, the signal decreases as increments the inhibitory capacity of the serum.

For detecting inhibitory antibodies against RBD Delta variant, an in-house sVNT was developed following the same design but using RBD of Delta variant as coating protein. Samples, positive and negative controls were diluted 1:25, followed by two serial dilutions 1/3 or 1/5 with PBS 0.3% BSA, 0.05% Tween 20. The dilutions were combined with equal volume of ACE2-His in a dilution plate. Fifty microliters of each mixture of ACE2 and serum dilution were added to separate RBD-coated wells of a 96-well plate (ThermoScientific 442404) and incubated at 25°C for 1 h. Bound ACE2 was detected with the conjugate anti-His tag: HRP (SB 105327-MM02TH) and developed using OPD/H_2_O_2_ as substrate. Absorbance was read at 492 nm on an ELISA microplate reader. In both assays, binding inhibition was calculated as (1− OD value of sample/OD value of Negative control) x 100%. Inhibitory titer was defined as the dilution in which each animal serum inhibits 20% of the binding of ACE2 without any competitor, calculated by linear regression using GraphPad Prism.

### Assessment of cellular immune response by IFN-γ ELISPOT

IFN-γ ELISPOT assay was performed using a Mouse IFN-γ ELISpot antibody pair (Mabtech, Sweden). Splenocytes were isolated in RPMI culture medium (Gibco, US). Samples (three or five mice per group) were processed individualized, with the exception of the control groups (Placebos) which were processed as pooled samples of three randomly selected mice. Duplicates cultures (5×10^5^ and 1×10^5^ splenocytes per well) were settled at 37°C for 48 h, at 5% CO_2_, in a 96 well round-bottom plate with 10 µg/mL of N_351-365_ peptide or N protein, 10 µg/mL of concanavalin A (ConA), or medium. After, the whole content of this plate was transferred to an ELISPOT pre-coated plate and incubated at 37°C for 16-20 h, at 5% CO_2_. The incubation conditions for conjugated antibodies and following steps were done as recommended by the manufacturers. A stereoscopic microscope (AmScope SM-1TSZ, USA) coupled to a digital camera was used for spots count.

### Statistical analysis

For statistical analyses the GraphPad Prism version 5.00 statistical software (Graph-Pad Software, San Diego, CA, USA) was used. Antibody titers were transformed to log10 for a normal distribution. For the non sero-converting sera, an arbitrary titer of 1:50 was assigned for statistical processing. The One-way Anova test followed by a Tukey’s post-test was used as parametric tests for multiple group comparisons. In case of non-parametric multiple comparisons, the Kruskal Wallis test and Dunns post-tests was employed. A standard P value consideration was as follows, ns, p>0.05; *, p<0.05; **, p<0.01; ***, p<0.001.

## 3. Results

### The recombinant nucleocapsid protein from SARS-CoV-2 Delta variant interacts with the ODN-39M

Based on the The N protein was subjected to the aggregation process with different quantities of ODN-39M for obtaining the aggregation curve (Figure 1A). Maximal aggregation (precipitation) of the mixture was obtained between 4 µg and 16 µg of ODN-39M. The condition of 60 µg ODN-39M (ODN over-saturated condition) corresponding to a mass ratio 0.66:1 protein/ODN, was selected for further characterization since precipitation phenomenon did not take place.

**Figure 1.**
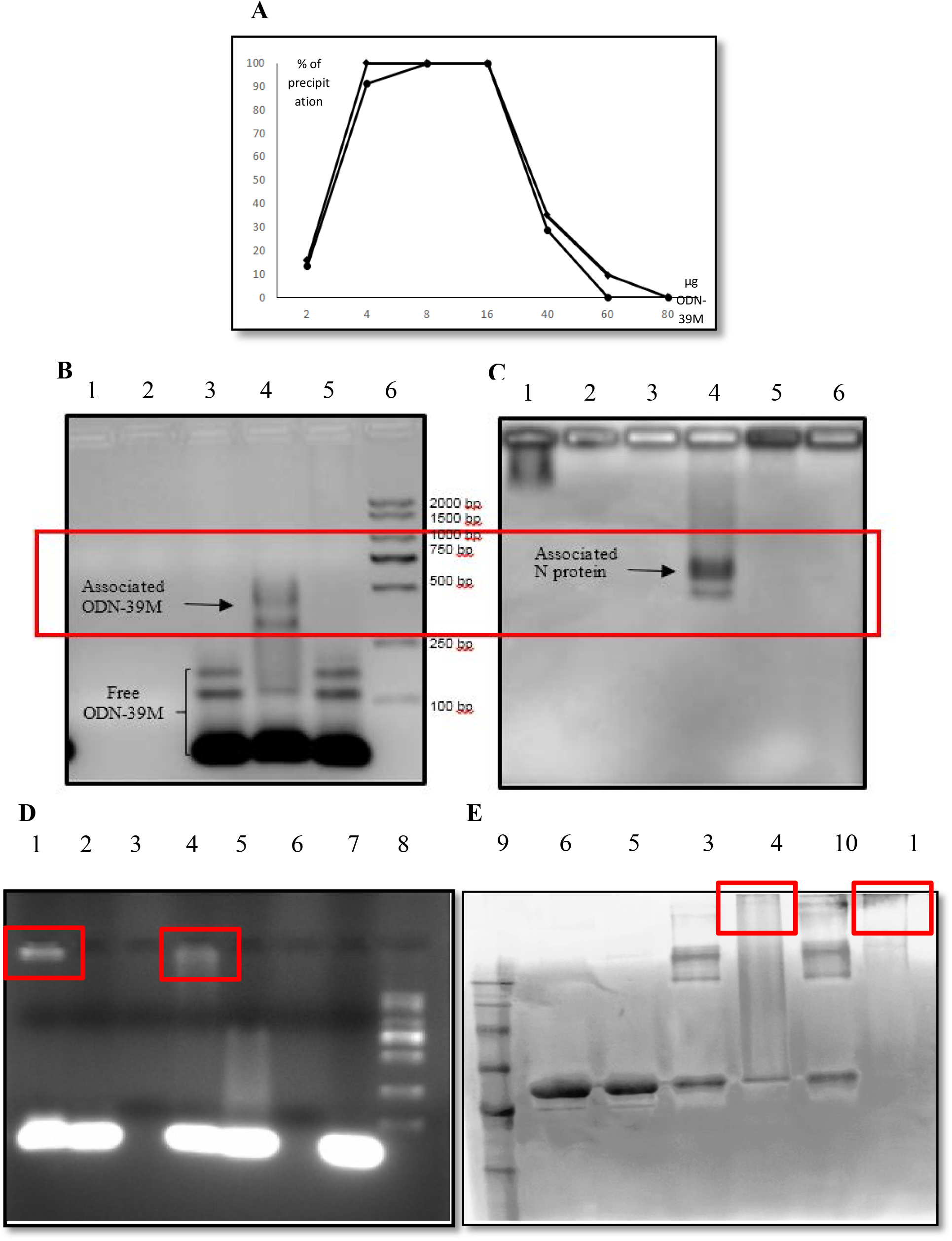
Characterization of the N+ODN-39 M preparation. A) Precipitation curve for different N:ODN ratios. Two independent experiments are represented. B and C) Analysis of the aggregation processes of N+ODN-39M, at 0.66:1 mass ratio (, by 2% Agarose gel electrophoresis stained with B) Ethidium Bromide and C) with Coomassie-Blue. 1: N protein; 2: Control protein; 3: ODN-39M; 4: N+ODN-39M; 5: Control protein+ODN-39M; 6: MW marker. Red box shows the zone where N protein and ODN-39M are co-localized. D and E) Analysis of different samples from the aggregation process of N+ODN-39M at 0.66:1 mass ratio after Formaldehyde (FA) treatment, by SDS-PAGE at 10%, both under non reducing conditions, stained with Ethidium Bromide and Coomassie-Blue E). 1: N+ODN-39M+1%FA; 2. ODN-39M+0.5%FA; 3. N+0.5%FA; 4. N+ODN-39M +0.5%FA; 5. N+ODN-39M; 6. N protein; 7. ODN-39M; 8. MW marker (DNA); 9. MW marker (protein); 10. N+1%FA. All FA reactions were quenched. Red boxes show the localization of both ODN-39M and N protein in each application point.

In order to have direct evidences of protein-ODN-39M interaction at the selected ratio, a sample of the preparation was applied into 2% agarose gel to visualize the co-localization of both molecules. The gel was stained with both, Ethidium Bromide to detect ODN-39M and Coomassie-Blue to detect the N protein. As shown in Figures 1 B and C, in the samples from the aggregation processes, a small fraction of the ODN-39M exhibited a delayed migration, in turn, the N protein migrated more compared to the protein without the ODN-39M. It means there was a common zone (in the indicated box) where the protein and ODN-39M were detected.

Finally, a sample of the N+ODN-39M preparation, previously immobilized with formaldehyde as a crosslinking agent, was also run in SDS-PAGE, under non-reducing conditions. The pattern obtained clearly evidenced the presence of high molecular weight N protein aggregates in N+ODN-39M preparation, since the sample treated with 1% FA was completely retained in the gel well. The same retention was observed for the ODN-39M when the cross-linked sample was analyzed at 2% Agarose gel (Figures 1 D and 1E).

### The N+ODN-39M combination, administered by intranasal route, is immunogenic in Balb/C mice

A first mice experiment was conducted to explore the immunogenicity of the N+ODN-39M combination administered by intranasal and subcutaneous routes. As shown in Figure 2A, all groups receiving the N protein by subcutaneous route induced high levels of anti-N antibodies in sera. In contrast, by intranasal route, only the group that received N+ODN-39M was able to induce a positive response, with titers similar to those obtained by subcutaneous route. The IgG1 and IgG2a subclasses against N Delta protein were also determined (Figures 2C and 2D). The IgG1 levels induced by G4 (N+ODN-39M, intranasally administered) were significantly lower (p<0.01) compared to those elicited by groups 1 and 2 (subcutaneously inoculated). On the contrary, the IgG2a levels were significantly higher in the G4 (p<0.001). Taken together, these results strongly suggest that a Th1 pattern was induced in mice receiving the formulation N+ODN-39M by intranasal route, as confirmed by the IgG1/IgG2a ratio = 1 for this group (Data not shown).

**Figure 2.**
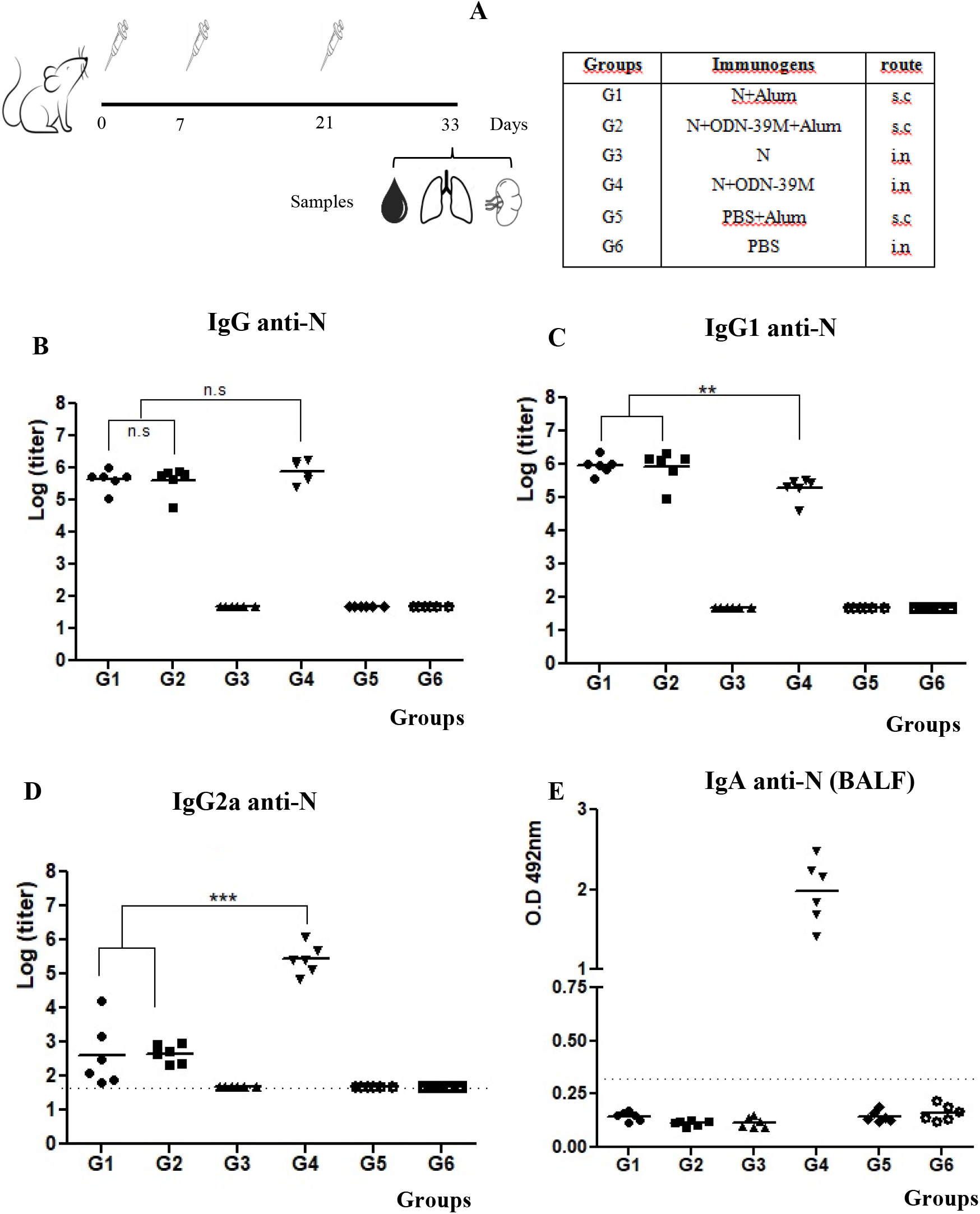
Evaluation of humoral immunity elicited by N formulations. A) Diagram of Balb/C mouse immunization. Six- to 8-week-old mice were immunized with three doses of each formulation according to the following group design: G1: N+Alum, sc. G2: N+ODN-39M+Alum, sc. G3: N (PBS), in. G4: N+ODN-39M, in. G5: N+ODN-39M, sc. G6: PBS+Alum, in. sc: subcutaneous, in: intranasal. Twelve days after the third immunization, mice were sacrificed and antibody responses in the serum and bronchoalveolar lavage fluid (BALF) were evaluated. B) C) and D) Systemic humoral immune response measured in sera by B) IgG ELISA against N, C) IgG1 ELISA against N and D) IgG2a ELISA against N. Data are represented as GMT of the titers. The statistical analysis was done by One Way Anova followed of Tukeys multiple comparison test. **p < 0.01, ***p < 0.001. E) Mucosal humoral immune response measured in BALFs by IgA ELISA against N. Data are expressed as O.D mean. The dotted line indicates the limit of positive response.

The IgA antibodies against N Delta were measured in BALF. As expected, only G4 (N+ODN-39M) elicited a positive response (Figure 2E). In turn, the anti-N IgG pattern of response found in BALF replicates the one observed in serum (data not shown).

To evaluate the CMI, the frequency of IFN-γ secreting spleen cells was evaluated after *in vitro* stimulation with two agents: the conserved peptide N_351-365_ and the N Delta protein. For the first stimulation agent two different determinations were done, at days 12 and 18, respectively (Figures 3A and 3B). At both times a similar pattern of CMI response was observed for all evaluated groups. A detectable IFN-γ secreting cell response was obtained in 3 out 3 animals only from G4, whereas no positive response was seen for animals from G3 (N protein in PBS, intranasal administered). In addition, only 1 out 3 mice elicited a positive response in each group immunized by subcutaneous route (G1 and G2). On the other hand, when the whole N protein was used as stimulating agent, the overall behavior was similar, however the number of responder animals in the groups subcutaneously immunized was increased to 3 out 3. Consistently with the previous determinations, the G4 (N+ODN-39M, intranasally administered) showed a clear trend to generate the highest response (Figure 3C).

**Figure 3.**
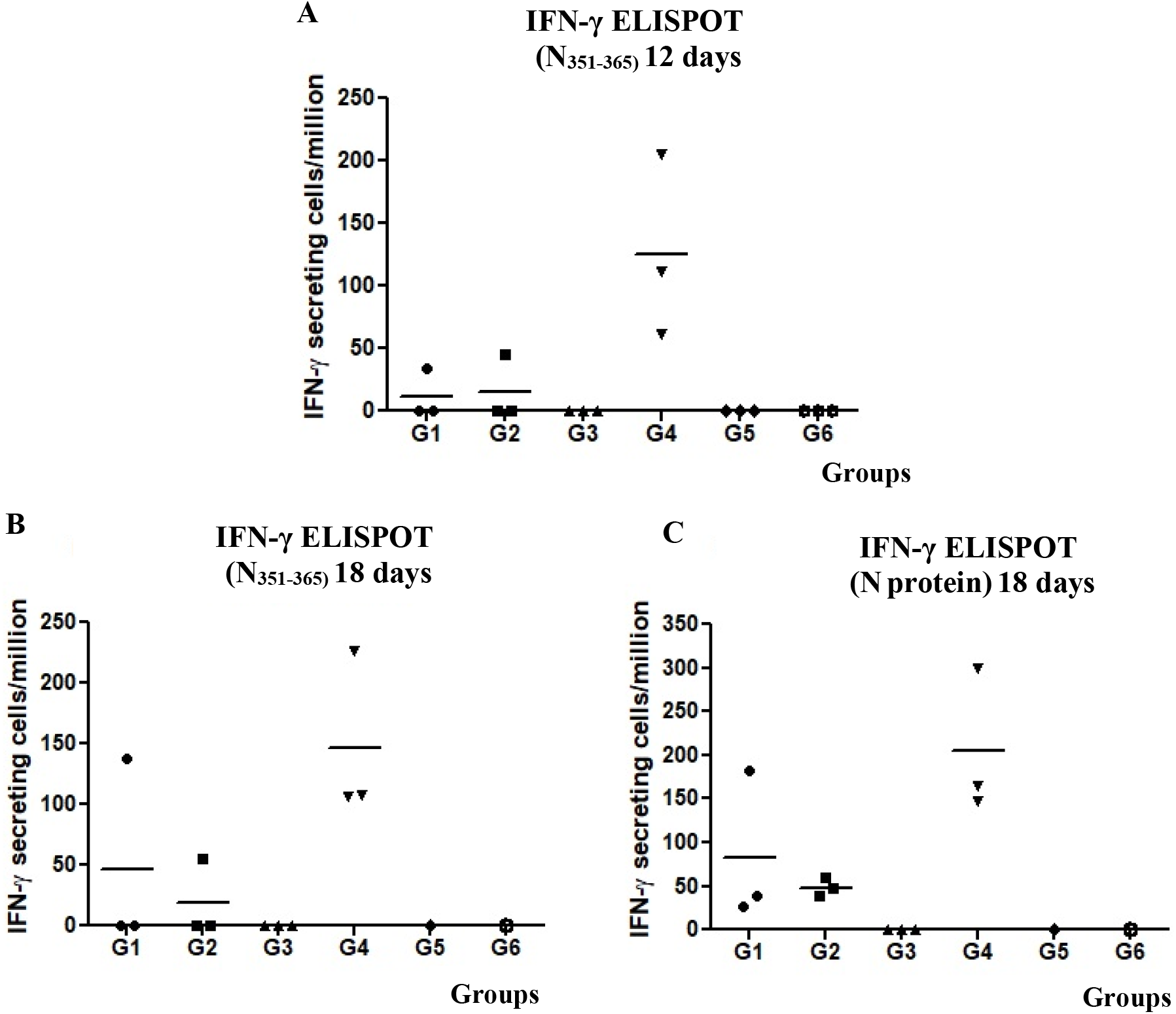
Cell-mediated immune responses induced in mice. Six- to 8-week-old mice were immunized with three doses of each formulation according to the following group design: G1: N+Alum, sc. G2: N+ODN-39M+Alum, sc. G3: N (PBS), in. G4: N+ODN-39M, in. G5: N+ODN-39M, sc. G6: PBS+Alum, sc. sc: subcutaneous, in: intranasal. Twelve days (A) or 18 days (B and C) after the third immunization, mice were sacrificed and spleens were extracted and *in vitro* stimulated with both N _351-365_ peptide (A and B) and N protein (C), and the frequency of the resulting IFN-γ secreting cells were measured by ELISPOT. The graphs show mean values (n = 3).

### Intranasally administered N+ODN-39M preparation induces a cross-reactive immune response, against N protein, until the Sarbecovirus level

Aiming at assessing the cross-reactive scope of the immunity generated by the combination N+ODN-39M administered by intranasal route, a second mice experiment was conducted. As shown in Figure 4A, the group inoculated with the combination N+ODN-39M induced in sera a positive IgG response against the N protein from SARS-CoV-2 Delta variant, SARS-CoV-2 Omicron variant and SARS-CoV-1. On the other hand, none of the groups showed a detectable response against N proteins from MERS-CoV and HCoV-229E (data not shown). A similar pattern of response was obtained when the IgA was measured in BALF samples against the different N proteins (Figure 4B).

**Figure 4.**
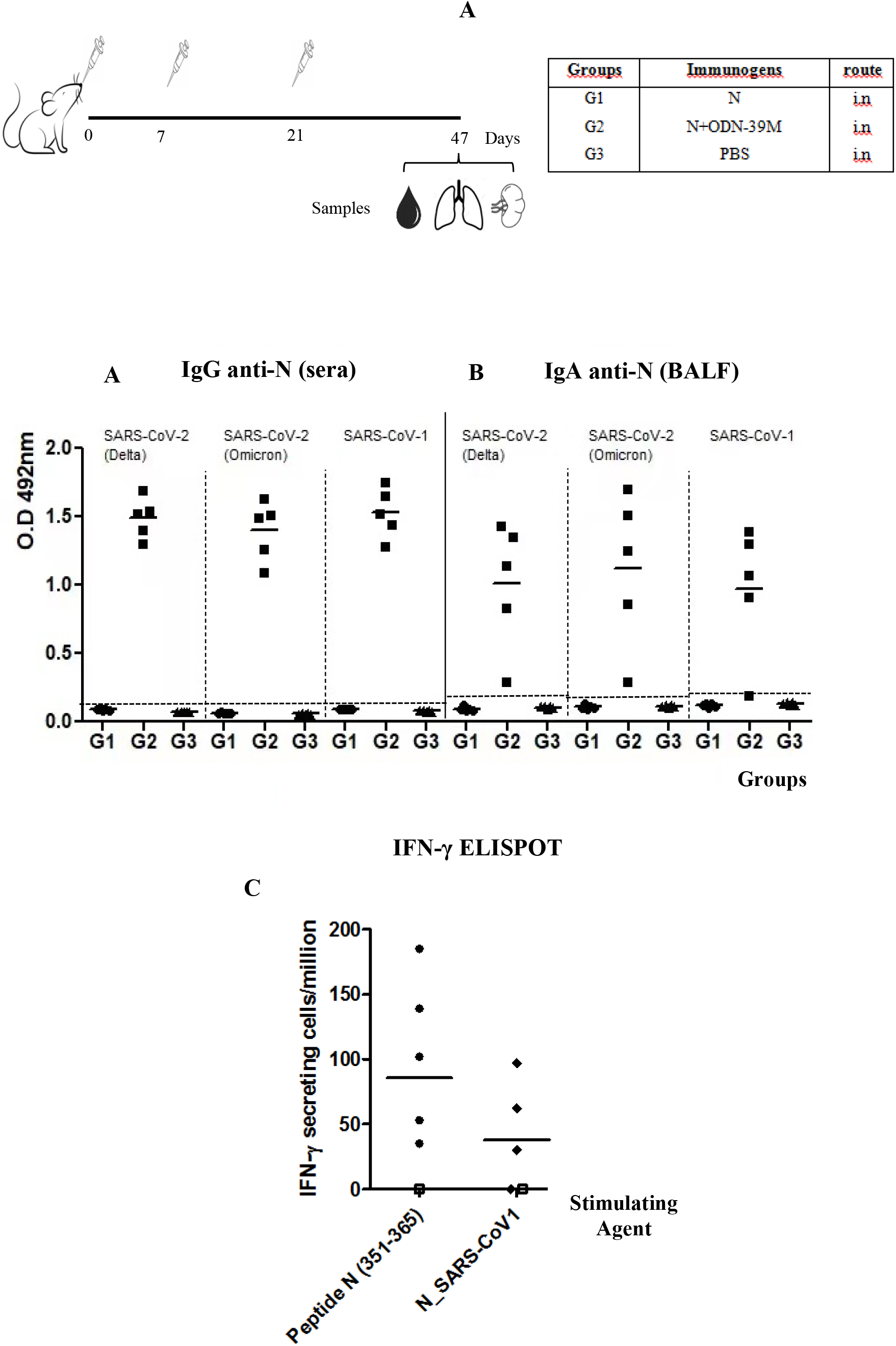
Cross-reactivity of the immune response generated by the intranasal administration of N+ODN-39M. A) Diagram of Balb/C mouse immunization. Six- to 8-week-old mice were immunized with three doses of each nasal formulation according to the following group design: G1: N protein, G2: N+ODN-39M and G3: PBS. Twenty-six days after the third immunization, mice were sacrificed, and antibody responses against N protein from SARS-CoV-2 Delta variant, SARS-CoV-2 Omicron variant and SARS-CoV-1 were measured by A) IgG ELISA in sera (n=5), 1:1000 dilution and B) IgA ELISA in bronchoalveolar lavage fluid (BALF), without dilution (n=5). Data are expressed as the mean of the O.D values. The dotted lines indicate the limit of positive detection. C) Cell-mediated immune response. Twenty-six days after the third immunization, mice were sacrificed and spleens were extracted and stimulated *in vitro* with both N_351-365_ peptide and N protein from SARS-CoV-1. The frequency of the resulting IFN-γ secreting cells were measured by ELISPOT. Square dots represent a pool of placebo samples. The graphs show mean values (n = 4 or 5).

To test the cross-reactive CMI, the frequency of IFN-γ secreting spleen cells was measured upon *in vitro* stimulation with: 1) the conserved peptide N_351-365,_ 2) N protein from SARS-CoV-1, 3) N protein from MERS-CoV, and 4) N protein from HCoV-229E. In line with the humoral immune response, animals receiving the intranasal administration of N+ODN-39M preparation exhibited a positive response against the peptide N_351-365_ and the N protein from SARS-CoV-1 (Figure 4C) whereas no response was detected for MERS-CoV and HCoV-229E proteins (data not shown). Results confirmed the cross-reactive nature of the CMI induced by N+ODN-39M preparation until Sarbecovirus subgenus level.

### The N+ODN-39M preparation exerts an adjuvant effect on RBD protein when both components are intranasally co-administered

A third mice experiment was conducted to explore the immunogenicity of a nasal bivalent formulation comprising N+ODN-39M+RBD. For anti-N response in sera, as shown in Figure 5B, G2 and G3 induced significant levels of Abs compared to the control group (p<0.001). The groups which received N protein without ODN-39M (G1 and G4) did not induce a positive response. The anti-N IgG levels elicited by G3 (bivalent formulation) were high (>10^3^), however statistically lower than those raised by G2 (p<0.001). In line with previous results, the groups G2 and G3 generated in sera a statistically similar anti-N IgG1, and IgG2a levels, suggesting a Th1 pattern (Figures 5D and 5F).

**Figure 5.**
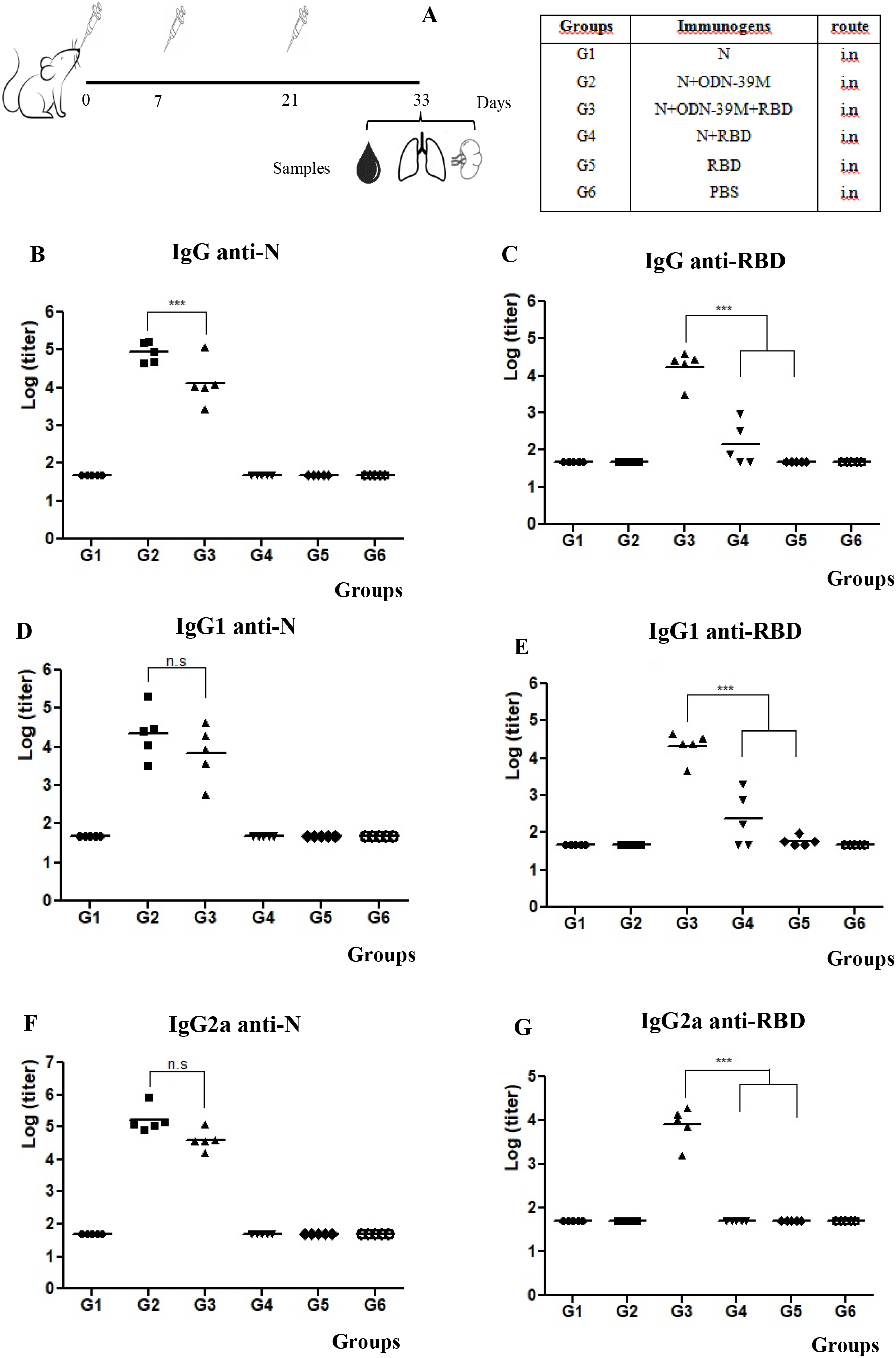
Pattern of the humoral immunity against N and RBD proteins in sera. A) Diagram of Balb/C mouse immunization. Six- to 8-week-old mice were immunized with three doses of each nasal formulation according to the following group design: G1: N protein, G2: N+ODN-39M, G3: N+ODN-39M+RBD, G4: N+RBD, G5: RBD, G6: PBS. Twelve days after the third immunization, mice were sacrificed, and antibody responses in the serum were evaluated by B) IgG ELISA against N, C) IgG ELISA against RBD, D) IgG1 ELISA against N, E) IgG1 ELISA against RBD, F) IgG2a ELISA against N, E) IgG2a ELISA against RBD. Data are expressed as the GMT of the titers (n = 5). The one-way Anova test followed by a Tukey’s post-test was used as parametric tests for multiple group comparisons. ***p <0.001.

Regarding the IgG antibody response in sera against RBD, only the group intranasally immunized with the bivalent formulation N+ODN-39M+RBD (G3) showed a 100% of seroconversion and IgG titers statistically higher than the rest (p<0.001) (Figure 5C). On the other hand, the group which received N+RBD (G4) showed a 60% of seroconversion (with titers ≤10^3^), and in the group immunized with RBD in PBS, none animal seroconverted. These results provide the first evidence of the adjuvant effect of the N+ODN-39M combination over RBD by intranasal route.

In turn, the pattern of IgG subclasses anti-RBD (Figures 5E and 5G) revealed higher IgG1 titers for G3 (bivalent formulation). In addition, only G3 was able to generate an IgG2a positive response when compared to the rest of the groups (p<0.001), suggesting again a modulation toward a Th1 pattern.

The assessment of IgA antibodies in BALF against both antigens is shown in Figures 6A and 6B. In both cases, only the groups inoculated with formulations containing the N+ODN-39M preparation elicited a positive response of IgA antibodies. Similar to the behavior observed for the anti-N IgG response generated in sera, the IgA response in BALF trended to be higher in the group receiving the monovalent formulation (G2) compared with the bivalent formulation (G3).

**Figure 6.**
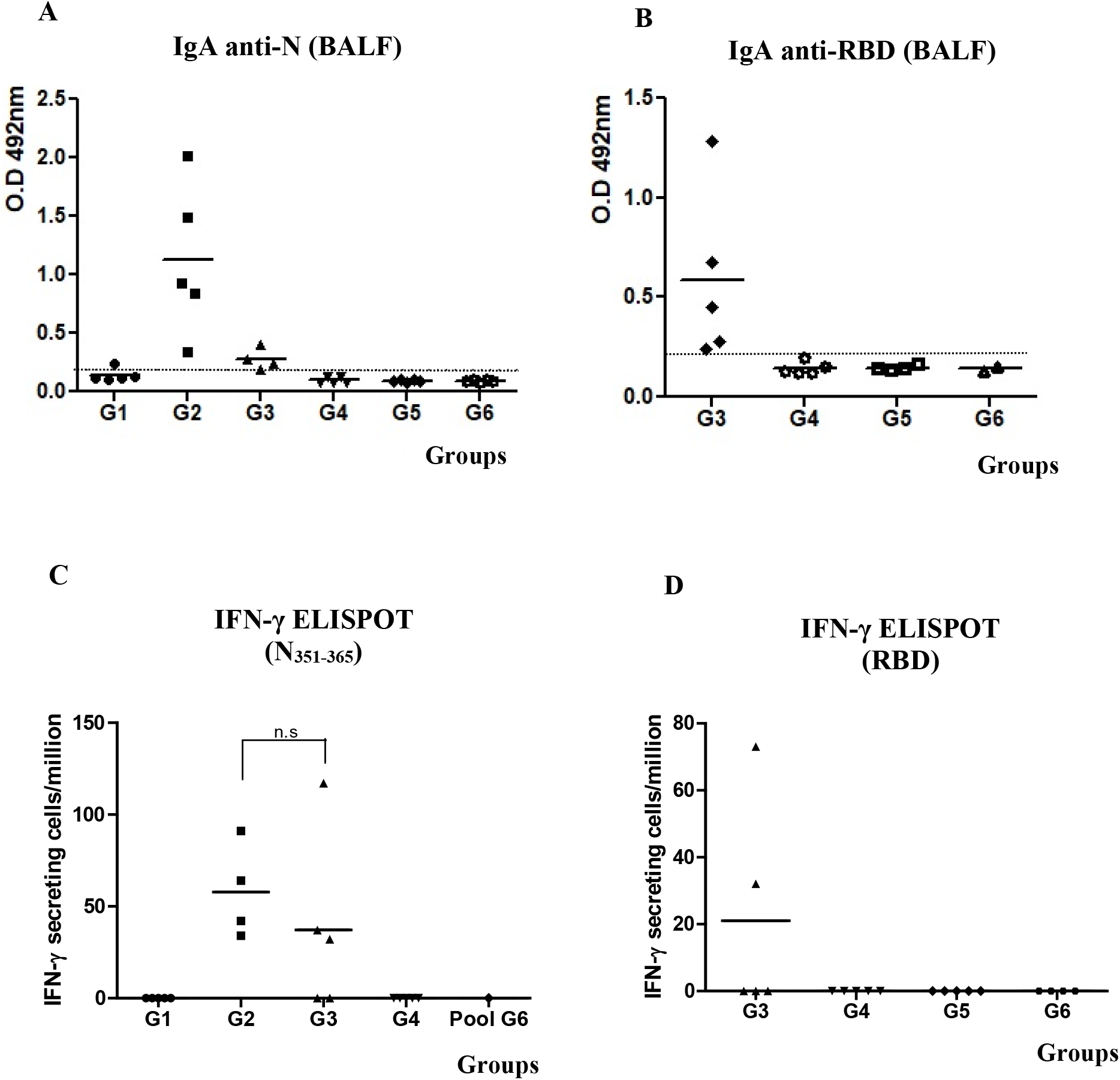
Evaluation of mucosal humoral immunity and cell mediated immunity for the bivalent nasal formulation. Six- to 8-week-old mice were immunized with three doses of each nasal formulation according to the following group design: G1: N protein, G2: N+ODN-39M, G3: N+ODN-39M+RBD, G4: N+RBD, G5: RBD, G6: PBS. Twelve days after the third immunization, mice were sacrificed, and antibody responses in BALFs (without dilution) were measured against A) N and B) RBD. Data are expressed as the mean of the D.O values. The dotted line indicates the limit of positive detection. The same day after the third immunization, for measuring cell-mediated immune response, spleens were extracted and stimulated *in vitro* with both N 351-365 peptide (C) and RBD (D). The frequency of the resulting IFN-γ secreting cells were measured by ELISPOT. Graphs represents mean values (n = 4 or 5). For statistical analysis the Kruskal Wallis test and Dunns post-tests was employed.

To explore the scope of the humoral immune response elicited by the bivalent formulation, sera and BALFs were also evaluated against the heterologous antigens: N from SARS-CoV-1 and RBD from Ancestral variant of SARS-CoV-2. Sera exhibited Ab titers against N from SARS-CoV-1 (Figure 7A) and RBD from Ancestral variant of SARS-CoV-2 (Figure 7B), following a pattern similar to that obtained for homologous antigens. On the other hand, in BALFs, 3 out 5 animals induced IgA Abs against RBD from Ancestral variant of SARS-CoV-2 whereas, all animals were positive against the homologous antigen (Figure 7C).

**Figure 7.**
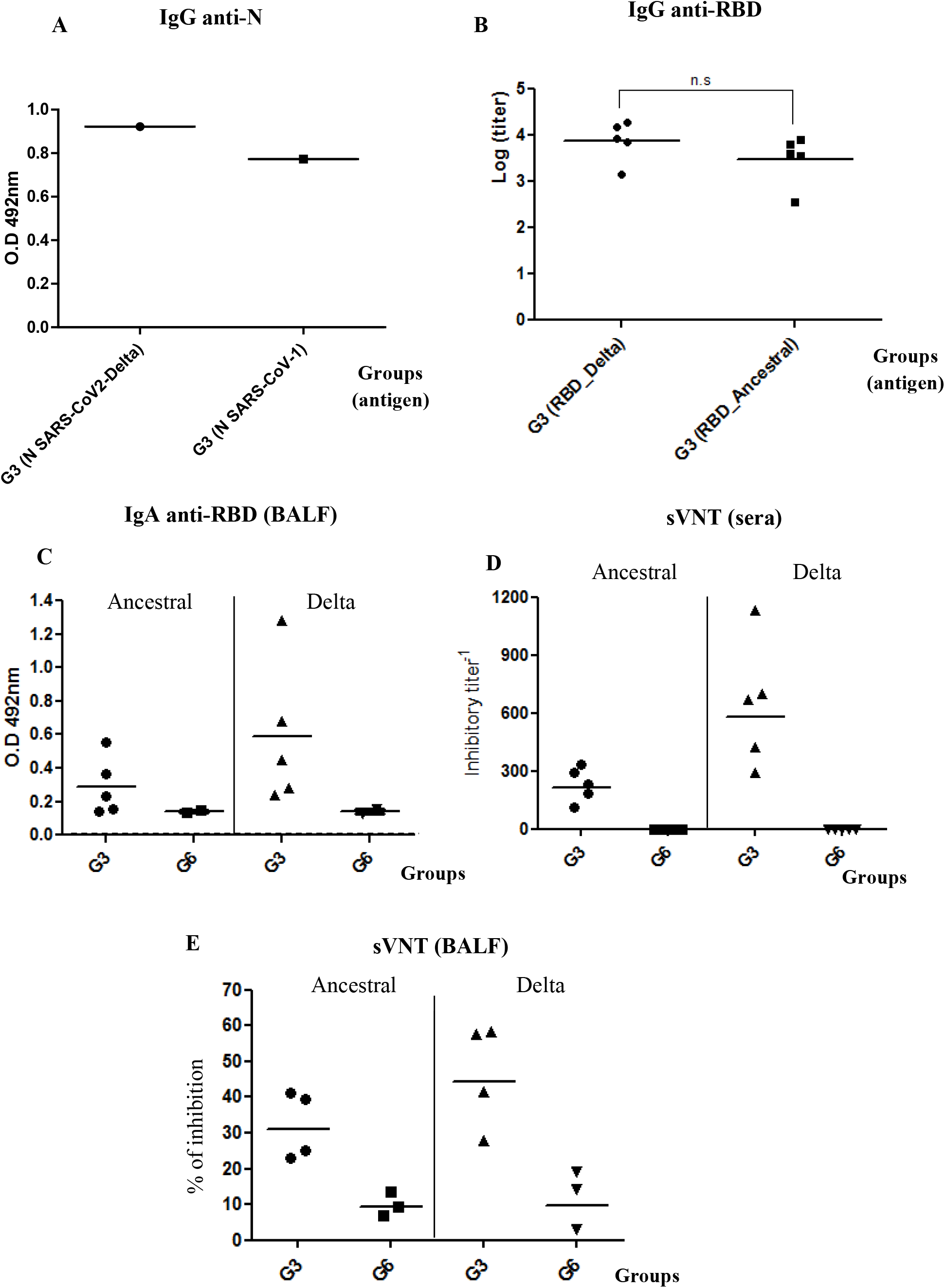
Evaluation of the cross-reactivity of humoral immunity induced by the bivalent formulation N+ODN-39M+RBD. Sera from groups G3 and G6 were evaluated by A) IgG ELISA against N proteins from SARS-CoV-2 Delta variant and SARS-CoV-1 (pool samples), B) IgG ELISA against RBD Ancestral and Delta variants. C) BALFs from groups G3 and G6 were evaluated by IgA ELISA against RBD Ancestral and Delta variants. Graphics for ELISAs represent either the mean of D.O values or GMT of titers. The dotted line indicates the limit of positive detection. D) sVNT against RBD Ancestral and Delta variants in sera and BALF, dilution 1:2 (E). Inhibitory titer was defined as the highest dilution in which each serum inhibits more than 20%.

In order to test the broad functionality of Abs elicited in sera and BALFs against RBD in the group receiving the bivalent formulation, a Surrogate Virus Neutralization Test (sVNT) was employed using the RBD from Delta and Ancestral variants of SARS-CoV-2. As shown in Figures 7D and 7E, all animal elicited antibodies with inhibitory activity against both variants. As expected, the higher values were obtained for the homologous RBD, and in sera samples.

Finally, results of CMI testing are shown in Figure 6C. Upon stimulation with the conserved N_351-365_ peptide, IFN-γ secreting cell response was statistically similar between G2 and G3 whereas no response was seen among splenocytes from the rest of the groups. In parallel, when splenocytes were stimulated with RBD Delta protein, a positive response was only obtained in 2 out of 5 animals from the group which received the bivalent formulation N+ODN-39M+RBD (Figure 6D).

## 4. Discussion

In the last two years, a number of studies have shown that N protein is a target of cellular immunity in humans and such a response has been correlated with protection against severity (Ferretti et al., 2020) (Kundu et al., 2022). Study published by Matchett *et al*., 2021, demonstrated that N protein, presented in the Ad5 platform, induced humoral and cellular immunity in mice, and such response correlated with protection against SARS-CoV-2 challenge (Matchett et al., 2021). Dangi et al, 2021, also evidenced that combining spike and nucleocapsid vaccine formulations improve distal protection in brain (Dangi et al., 2021). These authors even demonstrated the protective role of anti-N Abs, proven by passive transfer experiments (Dangi et al., 2022).

However, despite the encouraging results validating the N protein as an appealing component for a vaccine against SARS-CoV-2 (Dutta et al., 2020), two questions remain unanswered, 1) the breadth of the immunity generated and 2) its capacity to be immunogenic as recombinant protein, by intranasal administration using adjuvants suitable for human use.

The present work provides evidences to answer the two aforementioned questions. The N Delta variant, as a recombinant construct obtained in *E. coli*, was selected as antigen since Delta variant was the major circulating strain when these studies started. Given that the fundamental function of the SARS-CoV-2 N protein is to package the viral genome into a ribonucleic nucleoparticle, we hypothesized that the N protein could interact with the ODN-39M to form aggregate structures. The ODN-39M is a whole phosphodiester ODN containing CpG motifs, with a proven adjuvant capacity for other viral vaccine candidates (Suzarte et al., 2015, Olivera et al., 2020). Firstly, after incubation of N protein with ODN-39M at different proportions, a typical saturating curve was obtained. This behavior is consistent with the results published by Jack *et al*., 2021, where the N protein forms biomolecular condensates with viral genomic RNA both *in vitro* and in mammalian cells (Jack et al., 2021). Considering vaccine formulation requirements, we selected a point where N protein remains soluble after incubation with the ODN-39M. The analysis by agarose gel, stained with Ethidium Bromide to visualize the ODN and with Coomassie blue for detecting N protein, revealed a modification of the migration patterns for both molecules, as well as their detection in a common zone. It indicates that some level of interaction between these two molecules is taking place which constitutes the second evidence of interaction between them.

For the first immunological evaluation in mice, we explored the N+ODN-39M preparation, by two administration routes: intranasal and subcutaneous. Among the intranasal groups, only the group receiving N+ODN-39M elicited high levels of anti-N IgG Abs in sera, similarly to those elicited by the groups inoculated with the N in alum by subcutaneous route. In addition, only this group elicited anti-N IgA Abs in BALFs and the pattern of IgG subclasses revealed a typical Th1 response. This behavior is consistent with the results obtained in the CMI assay, where only the group immunized with N+ODN-39M by intranasal route, induced a positive response in all animals tested when the peptide N_351-365_ was used as stimulating agent. These experimental evidences highlight the key role of the N+ODN-39M preparation in the induction of a mucosal humoral anti-N response, as well as the CMI against the peptide N_351-363_. Of note, this peptide spans a conserved region among sarbecovirus which is immunodominant in SARS-CoV-2 Balb/C infected mice. Venezuelan equine encephalitis replicon particles vector (VRP), expressing this single SARS-CoV-2–specific CD4+ T cell epitope, partially protected mice from SARS-CoV-2 (Zhuang et al., 2021).

Three possible elements could contribute, at the same time, to the results obtained with the nasal N+ODN-39M preparation: 1) the conformation of the N protein in the mixture, 2) the adjuvant role of the ODN-39M and 3) the route of administration. The cross-linking experiments using formaldehyde clearly showed that after incubation of N+ODN-39M in the selected proportion, some changes took place in the protein structure, rendering higher molecular weight structures. A more detailed characterization of these aggregate preparations is currently ongoing. Such structures may contribute to the immunogenicity, since it is known that aggregate conformations are immunogenic for intranasal route (van Beek et al., 2021). On the other hand, CpG ODNs are considered very promising adjuvants (Bode et al., 2011). The ODN-39M evaluated in the present work is also a CpG ODN, but having a phosphodiester backbone, making the formulation N+ODN-39M very attractive for human use, since the thioate backbone has been associated to adverse reactions in therapeutics interventions (Bode et al., 2011). Based on the agarose gel profiles, we speculate that one portion of the ODN-39M interacts with N protein whereas another major portion remains free in the preparation. The ODN-39M associated to N protein, could be protected from a rapid degradation and its adjuvant capacity can be favored. Besides, the functionality of the free ODN-39M is not discarded, since it is known that other CpG ODN having a natural phosphodiester backbone also exerted adjuvant effect by intranasal route (Maeyama et al., 2014). In fact, mucosal tissues are enriched of plasmacytoid dendritic cells, which are easily activated by TLR9 agonists as the CpGs ODNs (Rothenfusser et al., 2002).

Based on the obtained results, the nasal N+ODN-39M preparation was selected for the second mice experiment to evaluate the breadth of the immunity induced. After measuring both arms of the immune system against the N protein of five coronaviruses, we observed that the immune response obtained reached the Sarbecovirus level represented by SARS-CoV-1. Such cross-reactive immune response was detected in the three kind of samples analyzed: BALFs, sera and spleens. Although the response obtained did not reach up to the level of Betacoronavirus genus where MERS-CoV is representative, we consider the cross-immunity reached is highly valuable. Several samplings for coronaviruses have been conducted in East and South East Asia, and around 50 SARS-related coronaviruses have been detected across 10 species of bat (Ravelomanantsoa et al., 2020). Accordingly, Crook JM *et al*, 2021, asserted that the highest the probability for homologous recombination of Sarbecoviruses through co-infection, the biggest the possibility of novel zoonotic emergence. (Crook et al., 2021).

The third animal experiment was addressed to assess the nasal bivalent formulation, composed by the N+ODN-39M preparation and the RBD from the Delta variant as inductor of neutralizing Abs. The recombinant RBD fragment, is a component of at least five approved vaccines for emerging use, such as the Cuban vaccines: Abdala and Soberana Series (01, 02 and Soberana Plus) (Hernández-Bernal et al., 2022), (Reed, 2022), and ZF001, a Chinese vaccine approved in China and Uzbekistan (Cao et al., 2022). Such vaccines have been capable of controlling the magnitude of the different infection’s waves of SARS-CoV-2 variants, nevertheless, as for the other approved vaccines, their capacity to halt the virus transmission has been limited. (Yu et al., 2020), (Wang et al., 2020).

In the present work, results demonstrated the adjuvant effect of the combination N+ODN-39M over the mucosal and systemic immune response generated by RBD in mice. The Ab response elicited in BALF and sera by the bivalent formulation recognized both, the RBD Ancestral and RBD Delta variants. Importantly, Abs in sera and BALF had inhibitory capacity against both RBD variants, and exhibited a Th1-like pattern. On the other hand, some animals exhibited CMI response in spleen against RBD. Taking together, we consider that the combination of N+ODN-39M constitutes a promising nasal vaccine component that can be added to the list of enhancers of RBD immune response by this route (Cao et al., 2021), (Du et al., 2021), (Jearanaiwitayakul et al., 2021), (Lazo et al., 2022), (Schild, 2021).

## 5. Conclusions

N+ODN-39M preparation, administered by intranasal route, is able to induce an anti-N cross-reactive immunity, at systemic and mucosal compartments, reaching Sarbecovirus level. In addition, it potentiates the immune response to RBD antigen, supporting its use as a potential component of a future intranasal pancorona vaccine. Particularly, the bivalent formulation N+ODN-39M+RBD constitutes a very promising vaccine candidate as a booster dose to amplify and broaden previous SARS-CoV-2 immunity generated by either, natural infection or vaccine, specially that one generated by inactivated vaccines.

## 6. Acknowledgments

We thank the colleagues from the Embassy of the Republic of Cuba in the People’s Republic of China, on behalf of the Cuban Government, for their direct contribution in the proper conduction of this project.

This work was supported by “National key R&D program of China (2021YFE0192200)”, “PNCT CITMA, Cuba”, “Hunan Provincial Base for Scientific and Technological Innovation Cooperation (2019CB1012)”, “The Science and Technology Innovation Program of Hunan Province, (2020RC5035)”, “Hunan Provincial Innovative Construction Program (2020WK2031)”

